# Exploring Parallel MPI Fault Tolerance Mechanisms for Phylogenetic Inference with RAxML-NG

**DOI:** 10.1101/2021.01.15.426773

**Authors:** Lukas Hübner, Alexey M. Kozlov, Demian Hespe, Peter Sanders, Alexandros Stamatakis

## Abstract

Phylogenetic trees are now routinely inferred on large scale HPC systems with thousands of cores as the parallel scalability of phylogenetic inference tools has improved over the past years to cope with the molecular data avalanche. Thus, the parallel fault tolerance of phylogenetic inference tools has become a relevant challenge. To this end, we explore parallel fault tolerance mechanisms and algorithms, the software modifications required, and the performance penalties induced via enabling parallel fault tolerance by example of RAxML-NG, the successor of the widely used RAxML tool for maximum likelihood based phylogenetic tree inference. We find that the slowdown induced by the necessary additional recovery mechanisms in RAxML-NG is on average 2%. The overall slowdown by using these recovery mechanisms in conjunction with a fault tolerant MPI implementation amounts to 8% on average for large empirical datasets. Via failure simulations, we show that RAxML-NG can successfully recover from multiple simultaneous failures, subsequent failures, failures during recovery, and failures during checkpointing. Recoveries are automatic and transparent to the user. The modified fault tolerant RAxML-NG code is available under GNU GPL at https://github.com/lukashuebner/ft-raxml-ng Contact: lukas.huebner@{kit.edu,h-its.org};, Supplementary information: Supplementary data are available at bioR*χ*iv.

## 1 Introduction

Failing hardware is projected to be one of the main challenges in future exascale systems (Shalf *et al*., 2011). In fact, one may expect that a hardware failure will occur in exascale-systems every 30 to 60 min (Cappello *et al*., 2014; Dongarra *et al*., 2015; Snir *et al*., 2014). High Performance Computing (HPC) systems can fail due to core hangs, kernel panics, file system errors, file server failures, corrupted memories or interconnects, network outages, air conditioning failures, or power halts (Gupta *et al*., 2017; Lu, 2013). Common metrics to characterize the resilience of hardware are the Mean Time Between Failure (MTBF) for repairable components and the Mean Time To Failure (MTTF) for non-repairable components. Both describe the expected average time for which a system will fully function *after* repair or replacement (Lu, 2013). For the sake of simplicity we will henceforth subsume MTBF and MTTF under the term MTTF and assume negligible repair and replacement time. Currently, most Message Passing Interface (MPI) implementations will terminate upon a rank failure. Regarding the failure frequency, we can therefore consider the entire set of MPI ranks as a number of serially connected single systems which will all fail if one single component fails. The MTTF of an MPI program running on Processing Elements (PEs) *n*_1_,*n*_2_,…,*n*_j_ with independent failure probabilities is there fore:

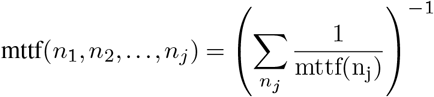

As the number of cores that can be used by scientific software increases, the MTTF decreases rapidly. Gupta *et al*. (2017) report the MTTF of four systems in the petaflops range containing up to 18688 nodes (see Table 1). Currently, most compute jobs only use a fraction of these petascale systems. Thus, current HPC jobs are not constantly aborted due to rank failures. In the not so distant future, on the by then commonly available exascale systems, scientific codes will run on hundreds to thousands of nodes and therefore experience a core failure every few hours (Cappello *et al*., 2014; Dongarra *et al*., 2015; Snir *et al*., 2014). We can therefore no longer ignore compute node or network failures, and require failure mitigating codes. To this end, we explore the current state of MPI failure detection technology and assess the amount of work required to make a massively parallel application fault tolerant by example of our RAxML-NG software (Kozlov *et al*., 2019), the successor of the widely used RAxML tool for phylogenetic inference (Stamatakis, 2014).

**Table 1.** MTTF of petascale systems as reported by Gupta *et al*. (2017)

| System | Nodes | Cores | MTTF |
| --- | --- | --- | --- |
| Jaguar XT4 (quad-core) | 7832 | 31328 | 36.91 h |
| Jaguar XT5 (8-core) | 18688 | 149504 | 22.67 h |
| Jaguar XT6 (32-core) | 18688 | 298592 | 8.93 h |
| Titan XK7 (16-core + GPU) | 18688 | 560640 | 14.51 h |

The remainder of this paper is structured as follows: In Section 2 we briefly discuss related work on fault tolerant scientific software in general and in Bioinformatics in particular. In Sections 3.1 and 3.2 we discuss the most common techniques and tools for implementing fault tolerance in parallel programs. We then describe the specific implementation in RAxML-NG in more detail in Sections 3.3 as well as 3.4 and present experimental results in Section 4. We conclude in Section 5 and discuss future directions of work.

## 2 Related Work

There already exist some fault-tolerant scientific applications. For example Ali *et al*. (2016) implemented a fault-tolerant numeric linear equation and partial equation solver. Obersteiner *et al*. (2017) extended a plasma simulation, Laguna *et al*. (2016) a molecular dynamics simulation, and Engelmann and Geist (2003) a Fast Fourier Transformation that gracefully handle hardware faults. Kohl *et al*. (2017) implemented a checkpoint-recovery system for a material science simulation. After a failure, the system initially assigns the work of the failed PEs to a single PE. The respective load-distribution algorithm (Schornbaum and Rüde, 2017) then recalculates the data distribution for the reduced number of PEs. Next, the PEs exchange the data residing on the wrong (i.e., overloaded) PE over the network via point-to-point communication.

In the field of Bioinformatics, some research into automatically restarting failed sub-jobs exists (Smith *et al*., 2006; Varghese *et al*., 2014). These methods require each job to be divisible into several sub-jobs and are not based on the checkpointing and restart paradigm. Finally, we are not aware of related work on assessing or implementing fault tolerance mechanisms in any other likelihood-based (i.e., ML or Bayesian inference) phylogenetic tree inference tool.

## 3 Fault Tolerance Techniques and Tools

Despite the currently limited support in MPI for mitigating hardware failures, some research on handling faults at the application level has been conducted. In the following we will outline our approach and the respective design rationales in comparison with other fault-tolerant scientific software.

### 3.1 Design Rationales and Implementation

The three main techniques for making programs fault tolerant are: Algorithm Based Fault Tolerance, restarting failed sub-jobs, and checkpointing/restart. Algorithm Based Fault Tolerance is used predominantly in numerical applications (Bosilca *et al*., 2008; Vijay and Mittal, 1997), but requires the algorithm in question to be extensible such as to include redundancy. We believe that it is not feasible to extend RAxML-NG in this way based on over 15 years of experience in RAxML-NG development.

Restarting failed sub-jobs as failure mitigation strategy is feasible when the program at hand can be split up into separate small and well-defined work packages which can easily be redistributed among nodes and managed via a (possibly distributed) work queue. RAxML-NG, however, applies a tightly coupled parallel iterative optimization strategy which cannot easily be split up this way, if at all. Finally, for deploying a checkpointing and restart strategy as presented here for RAxML-NG, the program has to regularly save its state to disk or memory.

Checkpoint/restart approaches can be further classified into system-level and application-level approaches. System-level approaches are (almost) transparent to the application, but are agnostic of the memory state subsets that can easily be recomputed at low computational cost (Hargrove and Duell, 2006; Roman, 2002). For instance, in any tree inference program that conducts phylogenetic likelihood calculations, the so-called Conditional Likelihood Vectors (CLVs, which store intermediate results of likelihood computations) dominate the memory requirements of the program and can require terabytes of memory in large-scale analyses (Jarvis *et al*., 2014; Misof *et al*., 2014). However, as these CLVs can be efficiently recomputed, they should not be saved in a checkpoint. We therefore chose to substantially extend the existing application-level checkpointing that was already implemented in RAxML-NG (see Section 3.3) to be more fine-granular as well as decentralized and therefore more scalable. Further, we use a fault-tolerant MPI implementation to make user interaction for restarts obsolete and lose substantially less work in case of failure. This removes a major hurdle for scaling to Exascale systems.

In general, we need to store the model parameters, branch lengths, and tree topology in a checkpoint. Typically, this data only needs a few megabytes or less and can be replicated in the local main memory of each rank. We can therefore store a full RAxML-NG checkpoint in the main memory of each rank. This approach is called diskless checkpointing and has the advantage of being substantially faster than writing checkpoints to disk (Plank *et al*., 1998).

In so-called coordinated checkpointing, all ranks of the program write their checkpoints at the same time which comes at the cost of additional synchronisation. Gavaskar and Subbarao recommend coordinated checkpointing for high-bandwidth, low-latency interconnections as they are common in modern HPC systems (Gavaskar and Subbarao, 2013). Since the ranks in RAxML-NG are synchronizing thousands of times per second anyway (Kozlov *et al*., 2015) we chose to conduct coordinated instead of uncoordinated checkpointing. The final design decision that needs to be taken, is whether to either use spare cores for replacing failed nodes or to simply reduce the number of nodes the job runs on upon failure. For example, Teranishi and Heroux (2014) describe a framework for recovering from failures by relying on available replacement processors. Making sufficient replacement processors available, however, would constitute a waste of resources in case there is no failure. Ashraf *et al*. (2018) study the performance implications of replacing failed nodes versus reducing the set of nodes. For their application they observed that shrinking represents a viable alternative to replacement. The overall CPU time required increased by a smaller degree when reducing the number of nodes. This is because the failed nodes were available for computations until the failure. We therefore redistribute the calculations to the remaining nodes upon failure in RAxML-NG.

We implemented fault tolerance for the most frequently used execution mode of RAxML-NG, that is, a phylogenetic tree search under maximum likelihood (ML). In addition, we focus on the fine-grained tightly coupled MPI parallelization scheme for supercomputers. Under this parallelization scheme, all sites of a large whole genome or multi-gene alignment are distributed via an appropriate load balancing algorithm (Kobert *et al*., 2014) to distinct MPI ranks, each running on a distinct physical core. In the following, we describe our implementation of the diskless checkpointing in Section 3.3 and the corresponding recovery mechanisms in Section 3.4. We call the resulting modified code FT-RAxML-NG (Fault Tolerant RAxML-NG).

### 3.2 The new MPI Standard and User Level Failure Mitigation

The upcoming MPI standard 4.0 will support mechanisms to mitigate failures of ranks or network components. Currently, there exist two actively developed MPI implementations which already support failure mitigation: MPI Chameleon (MPICH) (Gropp, 2002) starting with version 3.1 and User Level Failure Mitigation (ULFM) (Bland *et al*., 2013). We chose ULFM as the MPI implementation to develop a failure-mitigating version of RAxML-NG because the authors are also working on the standardization of MPI 4.0. We thereby hope to be as forward compatible as possible. Researchers have already used ULFM for other scientific software (Ali *et al*., 2016; Engelmann and Geist, 2003; Kohl *et al*., 2017; Laguna *et al*., 2016; Obersteiner *et al*., 2017). ULFM reports failures by returning an error on at least one rank which participated in the failed communication. This rank then propagates the failure notification to the other ranks via a dedicated MPI call. The next time a rank calls an MPI operation it will be notified that another rank revoked the communicator. Different ranks can therefore be in different parts of the code when they detect the failure. Next, all surviving ranks collectively create a new communicator excluding the failed ranks (Message Passing Interface Forum, 2017).

ULFM detects hardware failures via several mechanisms, depending, for example, on the interconnection type. One of the fundamental mechanisms to achieve this are regular heartbeat signals. If a rank misses three subsequent heartbeat signals, its observer will report that it failed. The MPI standard only mandates progress during MPI calls. For ULFM this means that, for the failure detection to work, at least one thread per rank has to enter an MPI function at regular intervals. If this is not the case, a rank cannot guarantee to respond to multiple consecutive heartbeat intervals. This would cause the rank to be falsely reported as being dead. We observed this behaviour when testing ULFM with the unmodified (i.e., non failure-mitigating) version of RAxML-NG in preliminary tests. All runs aborted because of false-positive failure reports. After consultation with the authors of ULFM, we adjusted some ULFM runtime settings. We increased the heartbeat interval, the heartbeat timeout, and enabled a separate dedicated heartbeat thread on each rank. Dedicated heartbeat threads ensure MPI progress at all times. This is because they are only responsible for sending heartbeats and can therefore enter the MPI runtime at all times without having to wait for the application to invoke an MPI function. In response to our discussion on the mailing list, the ULFM team published a tutorial on this topic on the ULFM website.1 After incorporating these changes into our configuration, we observed a substantial reduction of false-positive failure reports. In theory, these modifications come at the cost of performance. ULFM will require more time to detect failures because of the increased heartbeat timeout. The latency will also increase because multiple threads are now invoking MPI calls. However, in our implementation, the performance impact seems to be negligible.

### 3.3 Fine-grained Checkpointing

#### Modifications of Original Checkpointing in RAxML-NG

RAxML-NG already supports checkpointing. Thus, if a failure occurs, one can restart the program from the last checkpoint. However, these checkpoints are only written infrequently during the tree search and model parameter optimization procedures (i.e., several hours can pass between two checkpoints). To devise an efficient fault tolerant version of RAxML-NG we thus need to substantially increase the checkpointing frequency.

The tree search state of RAxML-NG consists of the model parameter values, the tree topology, and the branch lengths of the currently best tree (i.e., the tree with the currently highest likelihood score). The model parameters include the stationary frequencies, the nucleotide transition rates, and the rate heterogeneity model parameters. By design of the parallelization (Kozlov *et al*., 2015), the tree topology (including the branch lengths) being evaluated by the processors is identical and consistent across all ranks at all times. However, the currently *best* tree is not directly stored in memory. We therefore need to implement mechanisms to recover this currently best tree. Furthermore, large-scale phylogenetic analyses are typically partitioned, that is, different parts of the genome or individual genes are assumed to evolve under distinct models. Thus, for each partition, RAxML-NG estimates a separate, independent set of model parameters (stationary frequencies, transition rates, etc.). The model parameters of a partition are stored only at those ranks that have been assigned at least one Multiple Sequence Alignment (MSA) column of the respective partition. Therefore, the model parameters for a specific partition may only be saved by a single rank and can be lost if that rank fails.

#### Mini-Checkpointing

Our goal is to support failure mitigation for any set of ranks. Therefore, a simple solution consists in storing the model parameters of each partition at all ranks. Together with the copy of the current best tree topology (see below), we call this procedure “mini-checkpointing”. FT-RAxML-NG stores a copy of the most recent mini-checkpoint in the main memory of each rank. Each time a numerical optimization procedure (e.g., branch length optimization, substitution rates optimization, etc.) updates the model parameters of any partition, one dedicated rank per model broadcast the updated model. All other ranks then save these parameters, too. We experimentally assess the overhead induced by introducing mini-checkpoints for model parameters and mechanisms for restoring the tree search state in Section 4.2.

The model parameters of typical empirical datasets require a few MiB in size. Thus, we expect the number of partitions *m* to have a negligible impact on the runtime of a single broadcast. It does, however, have an impact on the number of broadcasts that FT-RAxML-NG needs to execute. The time required for checkpointing model parameters is thus in 𝒪 (min(*p, m*)· *T*_*bcast*_(*p*)), where *p* is the number of ranks. As already mentioned, we perform mini-checkpointing each time an optimization procedure updates the model parameters (see Figure 1). The mini-checkpoints are therefore also consistent across all ranks when we write a regular checkpoint to disk. Hence, we do not need to collect the model parameters for regular checkpointing as restoring them only constitutes a local operation that does not require any additional communication.

**Fig. 1.**
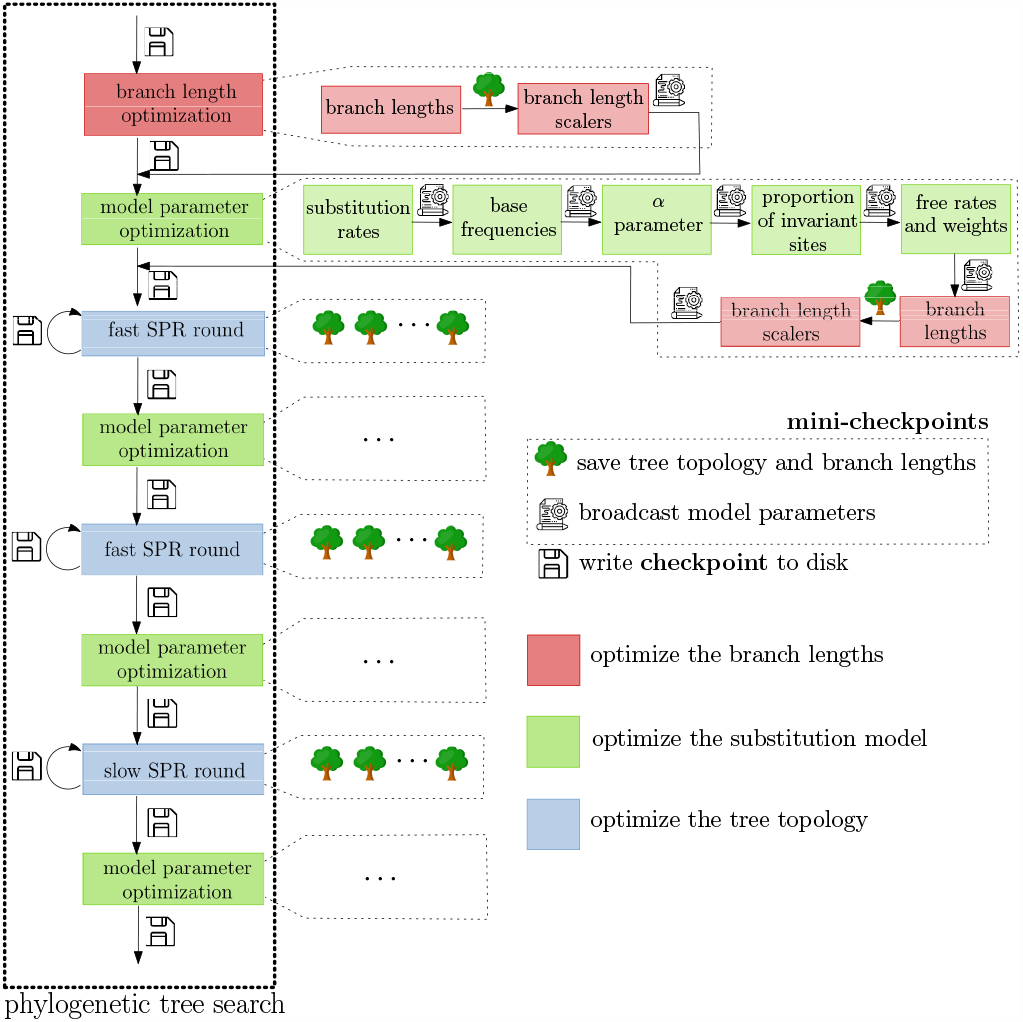
Frequency of checkpointing. On the left, an overview of the RAxML-NG tree search mode is given. RAxML-NG writes checkpoints to disk before each step of the optimization procedure and when optimization is completed. These are the regular to-disk-checkpoints which were already implemented. To obtain up-to-date model parameters, the master rank has to collect them first. Depending on the dataset and number of ranks used, each of these steps can take multiple hours to complete. By introducing mini-checkpointing, we increase the frequency at which model parameters are shared. The ranks now broadcast them after each sub-step, denoted by the respective model symbol. Additionally, the tree is saved each time it is changed, that is, after adjusting the branch lengths and during Subtree Pruning and Regrafting (SPR) rounds. Regular checkpoints are still written do disk, but do not need to collect and contain the model parameters, as these are already consistent across all ranks.

#### Saving the currently best tree topology

All changes to the tree topology are applied across all ranks simultaneously and consistently. Hence, we do not need to broadcast them. RAxML-NG does not explicitly store the currently best tree topology. A SPR round modifies this tree topology and saves the moves needed to restore the best tree topology in a rollback list. Modifying this rollback data structure to support restoring the best topology at the time of the last mini-checkpoint would be overly complex. To simplify recovery, we thus chose to copy the currently best tree to a separate roll-back data structure each time this currently best tree changes. As we do not need to broadcast the model parameters in this case, this merely constitutes a local operation on each rank.

### 3.4 Failure Recovery

If FT-RAxML-NG detects a failure during one of the model parameter optimization procedures or during an SPR-round that modifies the tree topology, it will restart the computation from the last mini-checkpoint. First, the surviving ranks need to agree on the ranks that are still alive and restore the search to a valid state. Evidently, as little computational work as possible should be lost because of a failure. FT-RAxML-NG thus needs to mitigate the failure, restore the search state, and resume the tree search without user intervention.

After ULFM has established a new communicator that contains only the surviving ranks, the search is still in an invalid state as no data has been redistributed or reloaded. Also, no rank is yet responsible for calculating the likelihood scores of the alignment sites assigned to the failed ranks. To simplify search state recovery, we implement this as a local operation, that is, no communication between ranks is required until search state restoration has completed.

To restart the tree search, we re-execute the load balancer (Kobert *et al*., 2014), reload the MSA data, and finally, restore the search state. The load balancer redistributes the MSA sites to the reduced set of ranks. Each rank then loads its respective MSA sites using partial loading. In partial loading, each rank selectively reads only those parts of the MSA file from disk that it requires for its fraction of the likelihood computations. To restore the search state, we locally copy over the model parameters from the last successful mini-checkpoint and the currently best tree from the local backup copy. Additionally, we need to invalidate and recompute internal caches, for example, the CLVs at the inner nodes of the tree.

### 3.5 Failures during checkpointing or recovery

If FT-RAxML-NG detects a failure during mini-checkpointing, it will restart from the preceding mini-checkpoint. Upon broadcasting of the model parameters, the ranks gather the received models in a temporary copy. All ranks then determine if a node failed during these broadcasts using `MPI_Comm_agree()`, which consists of three consecutive collective operations. If a failure occurred during checkpointing or the first two of these collective operations, a failure will be reported at all ranks. In this case, we discard the temporary checkpoint copies and start over from the preceding checkpoint that is still valid and stored in the working copy. If a failure occurs during the last collective operation which is part of `MPI_Comm_agree()`, all ranks defer reporting this failure to the next MPI call. At this point, all ranks are aware of the failure and can subsequently report it. As the first two collective operations of `MPI_Comm_agree()` did not re-port an error, the checkpoints stored in the temporary copies are valid and consistent across all ranks. Mini-checkpointing the tree topology is a local operation, if a rank fails during this step, the created mini-checkpoints is still valid on all other ranks. The surviving ranks can then restart the computation after detecting the failure.

The checkpoint-to-disk procedure will only fail if the master rank fails while writing the checkpoint to disk. In this case, the last checkpoint will still be valid and can be used to restart the search. If any other rank fails while the master rank writes the checkpoint, we will detect this failure at the beginning of the next optimization round. In this case, we can restart the computation from the checkpoint that was just written. As recovery is a local operation, it completes on each rank that does not crash. If additional ranks fail during the recovery, all surviving ranks will first complete the restoration process and subsequently handle the additional failure(s).

## 4 Results

### 4.1 Experimental Setup

We conduct all experiments on the ForHLR II supercomputer located at the Steinbruch Center for Computing (SCC) in Karlsruhe.2 It comprises 1178 worker nodes. Each node is equipped with two sockets of Intel Xeon E5-2660 v3 (Haswell) Deca-Core CPUs. All nodes are connected via an InfiniBand 4X EDR interconnection (Steinbruch Center for Computing (SCC), 2020). The files are served by two file server nodes from a DDN ES7K RAID with 14 volumes.

We use OpenMPI v4.0, ULFM v4.0.2u1, and GCC 9.2 for our experiments where not mentioned otherwise. FT-RAxML-NG is based upon RAxML-NG `c2af275ae6` on branch `coarse` released on March 5, 2020. See on-line supplement for more details on the hardware and software setup as well as the RAxML-NG/FT-RAxML-NG invocations.

#### Datasets Used

For our experiments, we use empirical protein (amino acid (AA)) and desoxyribonucleic acid (DNA) datasets with a varying number of taxa (36 up to 815 species/sequences), alignment lengths (20364 up to 21410970 sites), and partition counts (1 to 4116) (see the online supplement for a description; all datasets are are available on-line3).

#### Failure Simulation

We simulate failures by sending the SIGKILL signal to the respective MPI processes when using ULFM v4.0.2u1 and by splitting the communicator when using OpenMPI v4.0 (see on-line supplement for options and a respective rationale).

### 4.2 Experimental Results

#### ULFM Overhead

We determined the impact of the heartbeat timeout and heartbeat thread settings on the failure recovery time of ULFM by repeatedly simulating failures and measuring the time until a new communicator is created. (Figure 2).

**Fig. 2.**
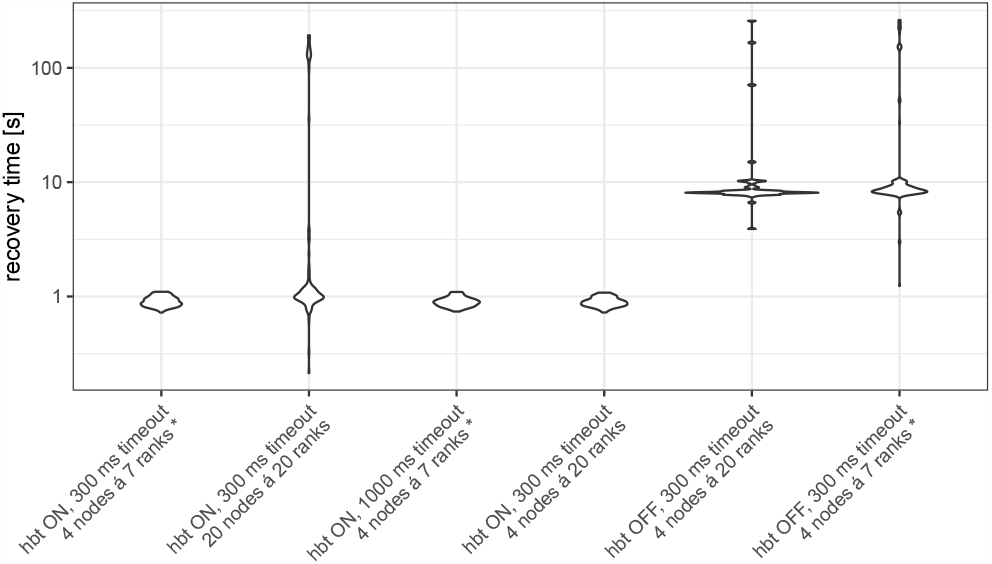
Violin plot of the time required by ULFM to detect and recover from a node failure. This includes the time for for all ranks to agree on which nodes have failed (handled by ULFM) and creating the new communicator. We measured two heart-beat timeouts: 300 ms (default, false-positives) and 1000 ms (no false-positives). Measurements marked with a * had additional nodes allocated which our code did not use, leaving them free for ULFM. The hbt (heartbeat thread) symbol indicates whether we enabled a thread responsible solely for sending heartbeat signals. Each measurement was performed at least 49 times.

First, we measured the time required for failure recovery under the default configuration (300 ms no heartbeat thread) on 4 nodes. Enabling the heartbeat thread decreased detection time from 8 s (median) to 900 ms (median). As the heart-beat thread also decreases the probability of false-positive failures, we therefore decided to keep this setting for all subsequent experiments. Next, we investigated the impact on failure detection speed when setting the timeout to 1000 ms. Contrary to our expectations, increasing the heartbeat interval did not change the result significantly (effect below standard deviation). We also investigated if slow failure detection is related to the computational load on the cores. For this, we executed three runs with different heartbeat thread and timeout settings. In these experiments we did not use all available cores for phylogenetic likelihood computations to exclusively reserve some for the ULFM runtime. This did however not change the ULFM failure recovery times. We also performed one experiment with 400 ranks to assess failure detection scalability. With the heartbeat thread enabled and a 300 ms timeout, 11.4 % of the recoveries required more than 2 s and 8.9 % of the recoveries required more than 90 s. These results are probably highly dependant on the specific HPC system used and should therefore not be generalized. Laguna *et al*. (2016) reported that ULFM required 11 s to recover when using 260 ranks on another system. These delays are induced by the underlying fault tolerant MPI implementations. When more optimized fault tolerant MPI implementations become available, FT-RAxML-NG will recover from failures faster.

To evaluate the slowdown induced by using a fault tolerant MPI implementation, we also compare the runtimes of *unmodified* RAxML-NG using ULFM with heartbeat threads enabled and disabled vs OpenMPI v4.0 as a baseline on three different datasets and PE counts (Table 2). We chose Open-MPI v4.0 as reference because ULFM v4.0.2u1 is based upon OpenMPI v4.0. The slowdown induced by ULFM ranges between *−*0.6 and 8.6 % of the reference runtime. The additional slowdown induced by separate heartbeat threads ranges between −0.7 and 2.2 % of the reference runtime. The additional slowdown induced by using a separate heartbeat thread compared to not using one ranges between −0.7 and 2.2 %-points of the reference runtime. The ULFM run that is faster than the OpenMPI reference run might be due to measurement fluctuations, but we did not further investigate this.

**Table 2.** Influence of ULFM on the runtime of unmodified RAxML-NG. We show the runtime using OpenMPI v4.0 and ULFM v4.0.2u1 with heartbeat thread (hbt, a dedicated thread for sending heartbeat signals) enabled and disabled. The slowdown is calculated relative to OpenMPI v4.0. #n: number of nodes, #r: number of ranks

| dataset | #n | #r | OpenMPI<br>[s] | slowdown [%] |  |
| --- | --- | --- | --- | --- | --- |
|  |  |  |  | ULFM<br>hbt: ON | ULFM<br>hbt: OFF |
| ChenA4 | 8 | 160 | 5582 | 0.1 | $-0.6$ |
| SoltD10 | 18 | 360 | 1437 | 1.9 | 2.5 |
| ShiD9 | 1 | 20 | 20911 | 6.4 | 8.6 |

Finally, we observed that ULFM sometimes reported a single rank failure but ULFM’s `MPI_Comm_shrink()` did not return the same communicator on all ranks. Multiple ranks reported their rank id as being 0 and a world size of 1. We reported this behaviour on the ULFM mailing list and the authors of ULFM reproduced and confirmed the bug (Bosilca, 2020). There is no patched version available yet.4 For this reason we use OpenMPI v4.0 as default MPI implementation and simulate failures as described in the on-line supplement for all performance evaluation experiments where we need to simulate failures.

#### Overhead of mini-checkpointing

We also assess the slow-down induced by mini-checkpointing (model-parameter updates and updates of the best known tree) as a function of the number of models (partitions) and ranks used. As the number of model-parameter updates and tree updates performed during a tree search is not the same, we show the overall amount of time spent performing either of those two in relation to the overall runtime of the tree search itself (Figure 3). For all test datasets mini-checkpointing requires less than 0.3 % of overall runtime. In the tree searches we executed, RAxML-NG performed 29 to 5814 model parameter updates and 242 to 14203 updates of the currently best known tree.

**Fig. 3.**
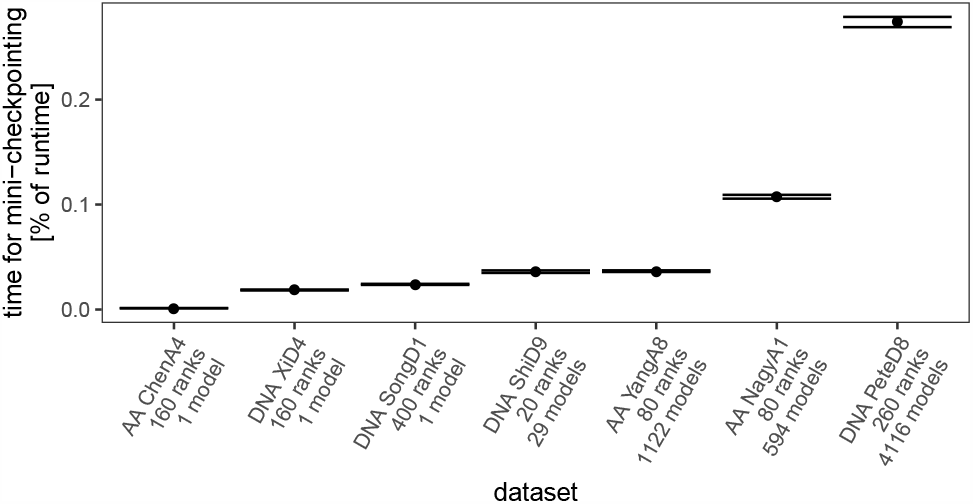
Proportion of overall runtime required for model parameter broadcasting and tree updates (“mini-checkpointing”). The mean (dots) and standard deviation (error bars) of the fraction of overall runtime required for mini-checkpointing across all ranks is shown. Each node has 20 CPU cores and executes 20 MPI ranks. The empirical datasets are described in the on-line supplement. Only if the number of models *and* the number of ranks increases, the time required for redistributing model parameters also increases.

#### Time Required for Restoring the Search State

We further profile the different phases of the recovery procedure in Figure 4. For this, we simulate failures as described in Section 4.1 and measure the time required for distinct phases of the recovery procedure. For each run with 20 ranks we simulate 19 rank failures; for each run with at least 40 ranks we simulate at least 39 rank failures.

**Fig. 4.**
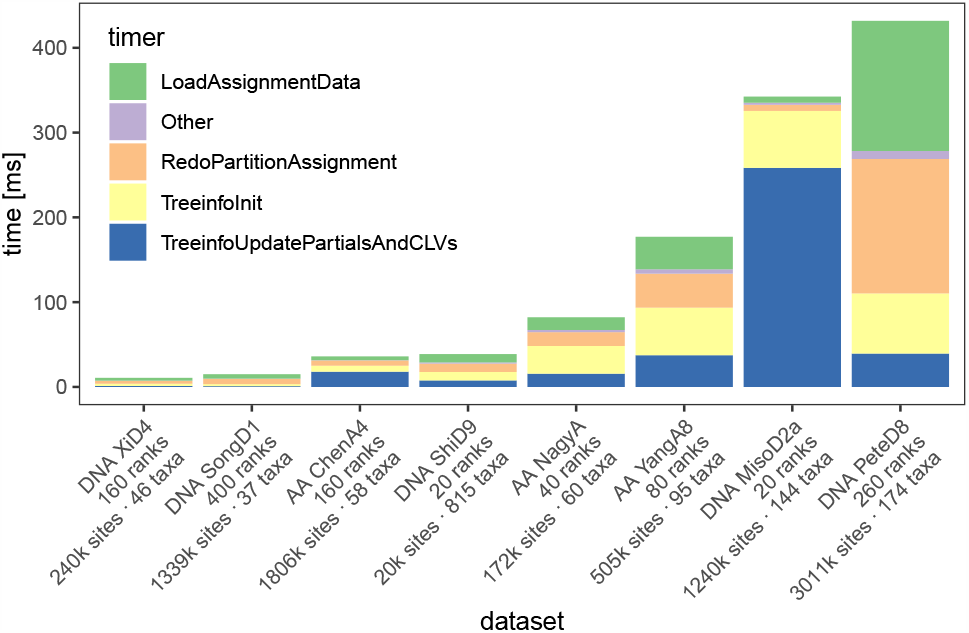
Time required for recovery from a checkpoint. “LoadAssignmentData” is the time required to load the MSA data from disk for which a rank is responsible. “RedoPartitionAssignment” is the time required for re-running the load balancer. “TreeinfoInit” is the time required for updating the `TreeInfo` data structure, including hundreds of memory allocations. “TreeinfoUpdatePartialsAndCLVs” is the time required for updating cached likelihoods. “Other” includes, for example, the time required for memory management and updating the data structures for mini-checkpointing.

We expect the time required for reloading the MSA data to increase with the product over the total number of sites and the number of taxa. On the ForHLR II, two file servers (see on-line supplement) handle disk accesses. This holds independently of the number of ranks used for the run. Thus, we expect the time for loading the MSA data to be independent of the number of ranks used. We further expect the time required for invalidating and recomputing the caches (e.g. CLVs) to increase with the number of sites allocated to each rank.

The time required for restoring the search state is below 100 ms in five out of eight runs, including one with 400 ranks. The remaining three runs are on datasets with more than 500000 sites as well as at least 95 taxa, and required up to 532 ms. The restoring times depend on the number of sites and the number of taxa. Considering DNA and AA datasets separately, the total restoring times required show a perfect rank correlation with the product of the number of taxa times the number of sites. That is, if a dataset *A* has more number of taxa times number of sites than a dataset *B, A* required more time to recover than *B*. The number of sites assigned to each rank directly influences the time required for updating the caches. For example, updating the CLVs of the MisoD2a dataset (see Section 4.1) which has over 62000 sites per rank required 258 ms.

#### Time Required for Reloading the MSA Data

We also investigate the time required to (re-)load MSA data from disk in a separate experiment as the MSA is (re-)loaded from a central disk array. Therefore, reloading these data can constitute a bottleneck. We measure the MSA loading time required for different datasets on ForHLR II. We distinguish between the initial (first time a rank loads data) load operation and further load operations (a rank loads data it has not previously loaded).

In each experiment, the load balancer decides which rank loads which part of the MSA. This is the initial load operation. Next, the load balancer shifts the data destination ranks by one, that is, rank 2 now loads the data of rank 1, rank 3 the data of rank 2 etc. We repeat this process until each rank has loaded each part of the MSA exactly once. We call all data loading operations after the initial one “further load operations”. Figure 5 shows the average time and standard deviation required to load a rank’s part of the MSA. The ForHLR II system has two separate file server nodes connected to the compute nodes via an EDR InfiniBand network.

**Fig. 5.**
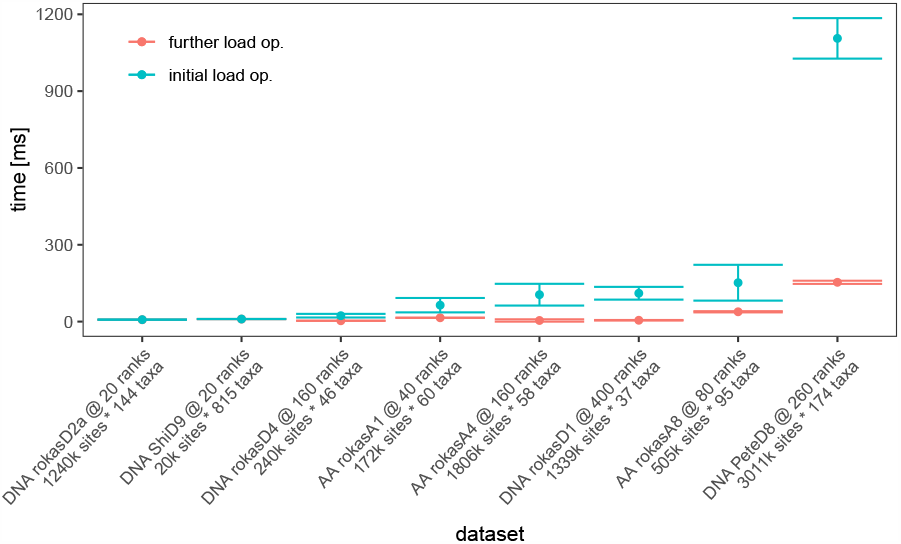
Time required for reloading the MSA from disk on the ForHLR II. Red dots and error bars (“initial load op.”) indicate the time required to load the rank’s part of the MSA the first time these data are accessed by any rank, that is, on program start-up. The blue dots and error bars (“further load op.”) represent the time it took a rank to load a part of the MSA it had never loaded before, but that had already been loaded by other ranks. These data can be assumed to be cached by the file servers.

The initial load operation represents an upper bound for the loading operation as the cache of the file system does not contain any copy of the data yet. The repeated loading of parts of the MSA, albeit from different ranks, is representative of the subsequent read performance as the filesystem cache already contains a copy of the data.

We provide the file sizes of the corresponding MSA files in the on-line supplement. For example, the DNA dataset PeteD8 has 3011000 sites and 174 taxa and a size of 500 MiB. This file size corresponds to the expected encoding size of 8 bit per site and per taxon.

In case of frequent failures, the cluster’s file system will cache the MSA data. This reduces the potential performance gain by an alternative approach that would store the data redundantly in the main memory of the compute nodes. If few failures occur, the MSA data will be accessed infrequently. In this case, the 1.2 s overhead for reloading the data can be amortized. As we performed these measurements on ForHLR II only, they can not be generalized.

With the goal of avoiding these disc accesses, we also implemented and tested a technique for tree-based compression of MSAs described by Ané and Sanderson (2005). This would allow to store a compressed copy of the entire MSA on each rank and could thus yield reloading the MSA from disk upon failure obsolete. Unfortunately, we found that the compression ratio (6*±* 7, not normally distributed) varies substantially across different empirical datasets. Thus, we conclude that this dedicated phylogeny-aware compression scheme is impractical for addressing the problem at hand (see on-line supplement for further details).

#### Overall Runtime Overhead

We measure the runtime overhead of FT-RAxML-NG running under OpenMPI v4.0 and ULFM v4.0.2u1 compared to unmodified RAxML-NG running under OpenMPI v4.0. If no failure occurs, that is the only additional work is the creation of mini-checkpoints, FTRAxML-NG is 2 *±* 2 % slower when using OpenMPI v4.0 and 8 *±*7 % slower when using ULFM v4.0.2u1. We also measure the runtime overhead for 10 (simulated) failures (see the on-line supplement). We use OpenMPI v4.0 for these measurements and do not reduce the number of compute ranks following simulated failures. This way we can measure the overhead induced by the recovery procedure. In our measurements, the slowdown of FT-RAxML-NG compared to unmodified RAxML-NG was 30 *±*20 % for 7–10 failures during one single run. In all experiments, the final log likelihood score deviated by less than 3*×* 10*−*8 from that of the failure-free reference run. A deviation in log likelihood scores might, for example, occur if there is a failure during a SPR round. In this case, we restart the SPR round from the currently best known tree topology and might therefore execute different SPR moves (see Kozlov *et al*. (2015)). You can find more details on these measurements in the on-line supplement.

## 5 Conclusion and Future Work

We present the, to the best of our knowledge, first study on fault tolerant parallel phylogenetic inference and one of the first studies on fault tolerance for Bioinformatics applications in general. It required approximately three person-months to design the fault tolerant version of RAxML-NG. We estimate that it would have required two person-months if the first author had been familiar with the RAxML-NG code beforehand. We demonstrate that our fault tolerant implementation can successfully handle and recover from multiple successive failures, including during critical parts of the program (e.g., during checkpointing and recovery). In addition, we provide a detailed study of the associated overheads for using a fault tolerant MPI version, executing the additional checkpointing mechanisms, re-loading and redistributing data from disk etc. With the goal of avoiding the disc-access when reloading the alignment data during recovery we implemented MSA compression. As the compression ratio was not as good as expected, we might explore other options to accelerate data reloading in the future. The program slowdown due to the additional checkpointing machinery amounts to 2% on average, while the overall slowdown including the deployment of ULFM amounts to 8% on average.

Given a sufficiently high frequency of job failures, we believe that a 2 % runtime overhead is alleviated by the corresponding savings in man hours and a shorter walltime-to-completion as manual job re-submissions are not required any more. The failure recovery time of but a few seconds is negligible assuming a MTTF of at least a few hours. Overall, the runtime overhead induced by checkpointing as well as the recovery times scale well with an increasing number of ranks. Nonetheless, the ULFM overhead of 6 % is high. As ULFM is still in its beta version, we are confident that this overhead will be reduced in future versions as it is critical for designing fault-tolerant parallel production-level applications. Future work will focus on implementing and assessing alternative data redistribution and redundant storage mechanisms.

## Supporting information

On-line supplement

## 6 Acknowledgements

Part of this work was funded by the Klaus Tschira foundation. This project has received funding from the European Research Council (ERC) under the European Union’s Horizon 2020 research and innovation program (grant agreement No. 882500).

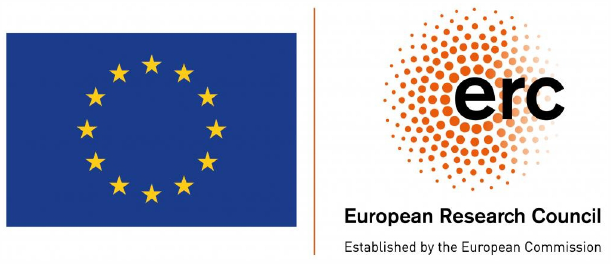

This work was supported by a grant from the Ministry of Science, Research and the Arts of Baden-Württemberg (Az: 33-7533.-9-10/20/2) to Peter Sanders and Alexandros Stamatakis.

## Footnotes

1 https://fault-tolerance.org/2020/01/21/spurious-errors-lack-of-mpi-progress-and-failuredetection/

2 https://www.scc.kit.edu/en/services/10835.php

3 https://figshare.com/s/6123932e0a43280095ef

4 As of 2020-11-30.

