## Supplementary material for "Exploring Parallel MPI Fault Tolerance Mechanisms for Phylogenetic Inference with RAxML-NG": On-line supplement

January 14, 2021

This is the on-line supplement to "Exploring Parallel MPI Fault Tolerance Mechanisms for Phylogenetic Inference with RAxML-NG". In Section 1 we describe the characteristics of the empirical datasets we use in our experiments. In Section 2 we describe the hardware, software, and tree search settings we use. In Section 3 we describe how we simulate rank failures when executing FT-RAxML-NG under ULFM and OpenMPI. In Sections 4 and 5 we discuss the impact of mini-checkpointing (without failures) as well as the combination of mini-checkpointing + recovery + additional work in case of (simulated) failures on the overall runtime. In Section 6 we describe our modifications to, and implementation of, the tree-based phylogeny-aware Multiple Sequence Alignment (MSA) compression scheme described by Ané and Sanderson [1].

#### 1 Datasets

Table 1 lists the characteristics of the empirical datasets used in our experiments.

**Table 1:** Characteristics of the datasets used for evaluating RAxML-ng and FT-RAxML-NG

| Designator | Data type | # taxa | # alignm. sites | # unique patterns | # partitions | file size [MiB] | Reference |
|---|---|---|---|---|---|---|---|
| SongD1 | DNA | 37 | 1,338,678 | 746,408 | 1 | 48 | Song *et al.* [15] |
| MisoD2a | DNA | 144 | 1,240,377 | 1,142,662 | 100 | 171 | Misof *et al.* [10] |
| MisoD2b | DNA | 144 | 413,459 | 371,434 | 50 | 57 | Misof *et al.* [10] |
| WickD3a | DNA | 103 | 436,077 | 422,676 | 14 | 43 | Wicket *et al.* [20] |
| WickD3b | DNA | 103 | 290,718 | 277,375 | 8 | 29 | Wicket *et al.* [20] |
| XiD4 | DNA | 46 | 239,763 | 165,781 | 1 | 11 | Xi *et al.* [21] |
| PrumD6 | DNA | 200 | 394,684 | 236,674 | 75 | 76 | Prum *et al.* [13] |
| TarvD7 | DNA | 36 | 21,410,970 | 8,520,738 | 1 | 736 | Tarver *et al.* [19] |
| PeteD8 | DNA | 174 | 3,011,099 | 2,248,590 | 4,116 | 500 | Peters *et al.* [12] |
| ShiD9 | DNA | 815 | 20,364 | 13,311 | 29 | 16 | Shi and Rabosky [14] |
| StamD10 | DNA | 436 | 1,371 | 1,011 | 1 | 0.6 | Stamatakis *et al.* [16] |
| NagyA1 | AA | 60 | 172,073 | 156,312 | 594 | 10 | Nagy *et al.* [11] |
| ChenA4 | AA | 58 | 1,806,035 | 1,547,914 | 1 | 100 | Chen *et al.* [2] |
| YangA8 | AA | 95 | 504,850 | 476,259 | 1,122 | 46 | Yang *et al.* [22] |

### 2 Experimental Setup

#### 2.1 Hardware and Software

We conduct all experiments on the ForHLR II supercomputer located at the Steinbruch Center for Computing (SCC) in Karlsruhe.[1] It comprises a total of 1,178 worker nodes. Each node is equipped with two sockets of Intel Xeon E5-2660 v3 (Haswell) Deca-Core CPUs with a clock rate of 2.1 GHz (max. 3.3 GHz) which results in a theoretical maximum peak performance of 832 GFLOPS per node. Each CPU has 64 KiB L1-cache (per-core), 264 KiB L2-cache (per core), 25 MiB L3-Cache (shared), and a 2,133 MHz bus as well as 64 GiB RAM. All nodes are connected via an InfiniBand 4X EDR interconnection [18]. Figure 1 shows the architecture of ForHLR II.

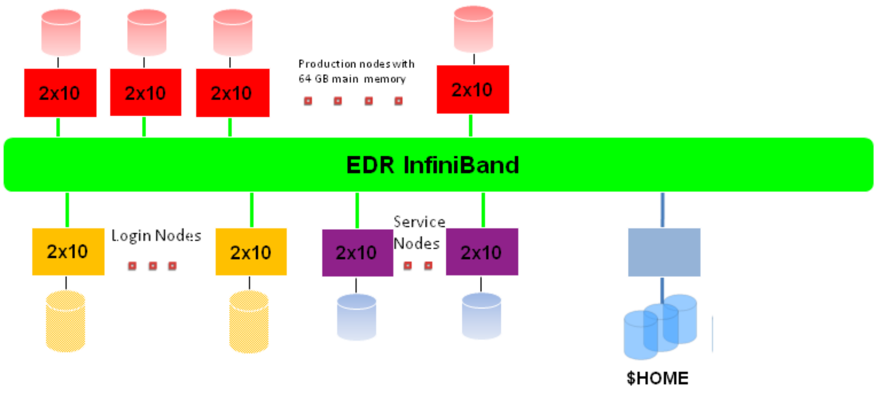


**Figure 1:** ForHLR II architecture. A worker node comprises a two socket system with 10 CPU cores each. All worker nodes and the file server nodes are connected using EDR InfiniBand. Image taken from the ForHLR II's website[1]; simplified to only show the part of the infrastructure we used.

ForHLR II has a Lustre distributed file system residing on a DDN ES7K RAID with 14 volumes. Each file is striped across 1 volume. We can read files from disk with a theoretical maximum I/O performance of $2\,\text{GiB}\,\text{s}^{-1}$ on a single node and $10\,\text{GiB}\,\text{s}^{-1}$ across all nodes. Two file server nodes provide file accesses [17]. Each of them has identical hardware as the compute nodes. The above is the default configuration on the ForHLR II. It is however possible to, for example, increase the number of stripes or file servers serving the files. We do not use this feature for our experiments as we intend to measure the performance for a typical use case, as typical RAxML-ng users will not manually tune their file-system configuration.

All nodes are running Red Hat Enterprise Linux (RHEL) 7.x and Slurm 20.02.3. We use OpenMPI 3.1 and ULFM v4.0.2u1 and GCC 9.2 for our experiments where not mentioned otherwise. FT-RAxML-NG is based upon `c2af275ae6` on branch `coarse` released on March 5th 2020.

#### 2.2 Tree Search Settings

We conduct the profiling experiments with the following RAxML-ng options when not stated otherwise. To use partial input file loading, we have to use random starting trees for initiating the tree search. The respective random seeds used are listed in Table 2. The following performance-relevant and model-specific RAxML-ng settings are used in each run: Tip-inner is disabled, pattern compression is enabled, per-rate scalers are disabled, site-repeats are enabled, the fast Subtree Pruning and Regrafting (SPR) radius is auto-detected, branch lengths scalers are proportional (ML estimate with NR-fast algorithm), the Single Instruction Multiple Data stream (SIMD) parallelization kernel is AVX2, and the number of threads per

[1] `https://www.scc.kit.edu/dienste/forhlr2.php`

Message Passing Interface (MPI) rank is one (see RAxML-ng[2] for details). The number of partitions (models) we use for each dataset is specified in each experiment.

**Table 2:** Random Seeds used in the Profiling Experiments

| dataset | random seed |
|---|---|
| NagyA1 | 1574547114 |
| ChenA4 | 1574484011 |
| YangA8 | 1574484011 |
| PeteD8 | 1574549152 |
| SongD1 | 1574549152 |
| MisoD2a | 1574443931 |
| XiD4 | 1574528895 |
| ShiD9 | 1574549152 |

### 3 Failure Simulation

We can simulate core failures in numerous ways without root access to the High Performance Computing (HPC) machines. When using User Level Failure Mitigation (ULFM) with no heartbeat thread, it suffices to put the application program into a long sleep to simulate a failure. When using a heartbeat thread, sending the `SIGKILL` signal to the process ID of a rank will simulate a failure. The program cannot catch, block ore ignore `SIGKILL`. It can therefore not perform any cleanup operation [8]. Other possible signals we can send include `SIGSEGV`, `SIGILL`, `SIGFPE`, `SIGBUS`, `SIGXFSZ`, `SIGPWR`, and `SIGXCPU`. None of these signals allow the receiving process to perform a cleanup operation. Our tests show that ULFM detects all of these simulated failures as rank failures with no noticeable difference, that is, the next MPI operation will fail with `MPI_ERROR_PROC_FAILED`. We are able to revoke the communicator using `MPI_Comm_revoke()` and the new communicator we subsequently build using ULFM's `MPI_Comm_shrink()` does not contain this "failed" node. We choose to simulate failures in experiments with ULFM by killing a process via signalling `SIGKILL` either by invocing `kill -SIGKILL` [6] or via self-signalling using `raise(SIGKILL)` [7].

When using OpenMPI we simulate failures either by calling `MPI_Comm_split()` such, that all ranks remain in the same communicator but the rank ids are shifted by one. This causes every rank to obtain a new site (data) assignment. We then proceed by executing a full recovery procedure. This allows us, for example, to measure the overhead induced by recovery but without having to account for the overhead caused by continuing with fewer ranks, after a rank failure.

### 4 Runtime Overhead Without Failures

We measure the runtime overhead caused by mini-checkpointing when no failures occur. We want to determine the runtime overhead caused by ULFM separately from the runtime overhead induced by our modifications to RAxML-ng. We therefore measure the runtime of FT-RAxML-NG with OpenMPI v4.0 and ULFM v4.0.2u1 as MPI implementations (see Table 3). In our measurements, the slowdown of FT-RAxML-NG running with OpenMPI v4.0 compared to the unmodified RAxML-ng running under OpenMPI v4.0 is $1.02 \pm 0.02$. The slowdown of FT-RAxML-NG running under ULFM v4.0.2u1 compared to the unmodified RAxML-ng running under OpenMPI v4.0 is $1.08 \pm 0.07$.

[2] `https://github.com/amkozlov/raxml-ng/wiki`

**Table 3:** Overall runtimes of unmodified RAxML-ng vs FT-RAxML-NG when no failures occur. That is, we perform mini-checkpoints (model updates) and tree updates but do not simulate failures. "s.dwn": slowdown

| type | dataset | ranks | OpenMPI RAxML [s] | OpenMPI FT-RAxML [s] | s.dwn | ULFM FT-RAxML [s] | s.dwn |
|---|---|---|---|---|---|---|---|
| AA | NagyA1 | 80 | 2,985 | 3,014 | 1.01 | 3,025 | 1.02 |
| AA | ChenA4 | 160 | 685 | 720 | 1.05 | 686 | 1.02 |
| AA | YangA8 | 80 | 1,182 | 1,230 | 1.04 | 1,210 | 1.02 |
| DNA | SongD1 | 400 | 1,365 | 1,383 | 1.01 | 1,541 | 1.13 |
| DNA | XiD4 | 160 | 3,760 | 3,858 | 1.03 | 4,466 | 1.19 |
| DNA | TarvD7 | 400 | 700 | 709 | 1.01 | 739 | 1.06 |
| DNA | PeteD8 | 260 | 5,393 | 5,492 | 1.02 | 6,197 | 1.15 |

### 5 Runtime Overhead With Failures

We measure the runtime overhead caused by failures (see Table 4). We use OpenMPI v4.0 for all measurements and simulate seven up to ten failures (see 4) as described in Section 3. The time points at which we simulate the failures are evenly distributed across the runtime of the runs. In our measurements, the slowdown of FT-RAxML-NG running with OpenMPI v4.0 compared to the unmodified RAxML-ng running under OpenMPI v4.0 is $1.3 \pm 0.2$. In all experiments, the final likelihood-score deviates by less than $3 \times 10^{-8}$ from that of the reference run. A difference in final log likelihood scores might, for example, occur if there is a failure during an SPR round. In this case, we restart the SPR round from the currently best known tree topology and therefore evaluate a different sequence of SPR moves [9].

**Table 4:** Overall runtimes of unmodified RAxML-ng ("reference") vs FT-RAxML-NG ("runtime") when failures occur. That is, we perform mini-checkpointing (models and tree updates) and simulate failures. We use OpenMPI v4.0 for all measurements and simulate failures as described in Section 3. "s.dwn": slowdown

| type | dataset | ranks | failures | reference[s] | runtime[s] | s.dwn |
|---|---|---|---|---|---|---|
| AA | NagyA1 | 80 | 10 | 2,985 | 3,077 | 1.03 |
| AA | ChenA4 | 160 | 10 | 685 | 1,093 | 1.60 |
| AA | YangA8 | 80 | 7 | 1,182 | 1,832 | 1.55 |
| DNA | SongD1 | 400 | 10 | 1,365 | 1,413 | 1.04 |
| DNA | XiD4 | 160 | 10 | 3,760 | 4,246 | 1.13 |
| DNA | TarvD7 | 400 | 10 | 700 | 998 | 1.43 |
| DNA | PeteD8 | 26 | 10 | 5,393 | 7,037 | 1.30 |

### 6 Tree-based Compression of Multiple Sequence Alignments

In this section, we describe our modifications and implementation of the tree-based phylogeny-aware MSA compression method described by Ané and Sanderson [1]. A more detailed description can be found in the following Master's thesis [4]. An Open Source implementation of the algorithm is available at `https://github.com/lukashuebner/ft-raxml-ng` on the "tree-based-msa-compression" branch.

An MSA consists of a set of aligned sequences. We will limit ourselves to Deoxyribonucleic Acid (DNA) sequences for now. This implies, that every sequence has the same length, possibly including gaps. Sequences can be, for example DNA or Amino Acid (AA). DNA has four states (`A`, `C`, `T`, `G`). In real world datasets, however, we sometimes want to encode that we are unsure of the exact nucleotide at a certain position. This is generally known as ambiguity encoding. An ambiguity can also mean that we observed multiple nucleotides at this position in different sequencing runs. The International Union of Pure and Applied Chemistry (IUPAC) defines 4 nucleotide states, 11 ambiguities codes and a gap [5] code. For example the character K represents either a G or a T at a position in the sequence. We expand the previously described [1] compression scheme to include this ambiguity encoding as it is required for analyzing empirical datasets and describe an algorithm to encode and decode the given MSA below.

The sequences of an MSA are located at the tips of a corresponding phylogenetic tree. The main idea of the tree-based compression scheme is to fully store only one sequence. We store all other sequences as a set of changes along the edges/branches of the tree (see Figure 2). We annotate these changes at the edges, for example $5 \to$ C means that the nucleotide state at the fifth site of the sequence changes to a C along the edge. This means that, ancestral states (inner nodes) have a sequence associated with them. For this compression approach, we evidently also need to store the corresponding tree topology for the specific compression as well. Using the most parsimonious tree (i.e., the tree that can explain the data by the least amount of mutations) guarantees the shortest encoding (best compression) [1].

For details on how to encode the tree in a binary Newick format and more details on how to encode changes along the branches of the tree, see Ané's and Sanderson's publication [1].

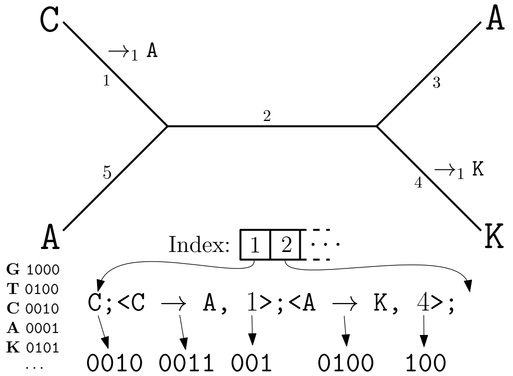


**Figure 2:** Encoding of the MSA. Encoding of a single site of the MSA. We encode the nucleotide's root state (upper left). Then, we encode the changes to the nucleotide state along the edges of the tree. We encode these changes in pre-order. We encode each change using the change mask and edge number (annotated next to the edges). We obtain the change mask by XORing the nucleotide states before and after the change. The index data structure points to the beginning of each site's encoding.

To encode MSA's sequences, we arbitrarily but deterministically choose one node as the root node. We thereby also defined define the order in which all nodes are visited during tree-traversal. We can thus encode the sequences in the MSA as a sequence at the root and the set of changes along the tree edges which result in the MSA's sequences at the tips of the tree. We then store the ancestral states of the sequence at the choosen root with four bits using a one-hot encoding. In a one-hot encoding, a single bit encodes for one of the four basic nucleotide states, that is, A = 0001, C = 0010, T = 0100, or G = 1000. A single nucleotide state is therefore 4 bits long. We encode ambiguities by setting multiple bits at once. We encode a gap by clearing all bits, thus distinguishing it from the state "any nucleotide" (N), which may also appear in the sequence. We store the changes to a site directly after the nucleotide state at this site at the root sequences. By doing so, we ensure that all the data we need for decoding the nucleotide states of one site are stored contiguously in memory, thereby enabling a cache-efficient decoding. Additionally, we do not need to store the index of the site this change modifies, which would need an additional log(sequenceLength) bits per change. We also store an index data structure $I$ mapping the site identifier to the start of the encoding for the specific site. We implement this via an array. We thereby implement random read access to the sites of the encoding.

We encode the changes of a nucleotide along the edges of the tree as the change's substitution mask and the edge this changes occurs on. We obtain the substitution mask by XORing the nucleotide state before and after the change. For example, if a T (0100) is replaced by a C (0010), the substitution mask will be

$mask = C \oplus T = 0100 \oplus 0010 = 0110$. This allows us to handle ambiguities.

We also identify the edges by a unique identifier. We obtain this identifier by enumerating the nodes in pre-order (parent, left child, right child) and assigning each edge the same id as the node it leads to. We chose pre-order instead of post-order as used by Ané and Sanderson [1] because this way we store the changes "root to tip". This yields the decoding more straightforward as we can implement it using a linear sweep without look-back through the encoding. We store a dummy entry in the index data structure pointing just past the last valid change to mark the end of the encoding.

#### 6.1 Description of the Algorithm

In this Section, we provide a description of the algorithm for compression and decompression of the MSA data. Compression consists of finding the ancestral states of the parsimony tree as well as encoding the changes along the tree. We refrain from encoding and decoding the tree itself and keep it in memory in an uncompressed format.

###### Computing the Ancestral States of the Parsimony Tree

We use Hartigan's [3] algorithm to calculate an assignment of sequences to inner nodes. This assignment has the property, that the number of mutations across the tree is minimal. It is not necessarily the only such assignment.

The algorithm takes a phylogenetic tree with fixed topology and fixed sequences at its tips as input. The algorithm consists of two phases (see Algorithm 1). The first phase assigns a set of possible ancestral states to each inner node of the phylogenetic tree. The second phase then selects one ancestral state per inner node. It does this in a way that minimizes the number of mutations across the tree.

**Algorithm 1** Hartigan's [3] algorithm: Overview

**Given:** A Tree $T$ with $i$ tips and the corresponding MSA $S$ with $i$ sequences $S_0 \dots S_i$

```
for each s_i ∈ S do
    Phase1(s_i, T)
    Phase2(s_i, T)
end for
```

To compute the possible ancestral states, we visit at each inner node in post-order (left child, right child, parent). This means that once we arrive at a node, we have already processed both children. For each tip, the set of possible states consists of the single fixed state in the input data. For all inner nodes, we check if the current node's children have possible common states. If they do, the current node's possible states are given by the intersection of the children's states. In case they do not, we assign the union of the children's possible states to the current node (see Figure 3.a and Algorithm 2).

To select an ancestral state for each node, we start at the root and traverse the tree in pre-order. This means, that we once we visit a node, we have already processed its parent. For the root, we choose one of the possible states at random. For each inner node, we choose its parents state, if this state is a possible state of the current node. If it is not, we choose a random state of the current node's possible states. In this case, one mutation occurred. For the tips, the states are already given, and we do not alter them. If the tip's state differs from its parent's state, a mutation occurred (see Figure 3.b and Algorithm 3).

In a bifurcating tree with vertex set $V$ and edge set $E$, $|E| = |V| - 1$ holds. Therefore, we can perform a Depth First Search (DFS) in $\mathcal{O}(|V|)$ time. As the sequences of the MSA are located at the tips, the tree has $|V| = 2n - 1$ nodes, where $n$ is the number of sequences in the MSA. In Phase 1, we compute a set intersection and possibly a set union for each node. There are only 16 possible values in the sets. We can therefore compute unions and intersections in $\mathcal{O}(1)$ time. We can use, for example, a binary set representation and bitwise `OR` and `AND` operations for this. In phase 2, we have to compute an element-of and random choice. We can compute element-of in $\mathcal{O}(1)$ time using a bitmask. We replace the random choice

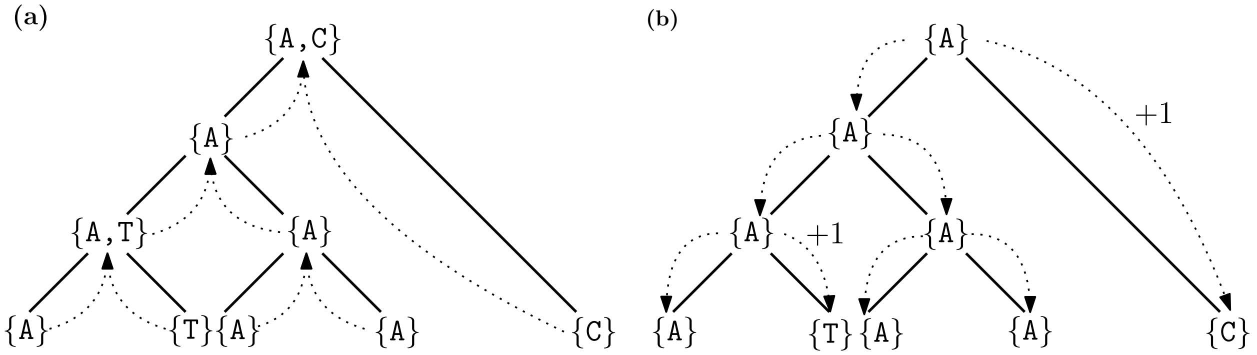


**Figure 3:** Reconstruction of the ancestral states as described by Hartigan [3]. A bifurcating phylogenetic tree with fixed topology and sequences at the tips is given. (a) In the first phase, we build possible ancestral states. If both children of the same parent have common states, we set these as possible ancestral states. If the two children do not have common states, we set the union of the children's states as possible ancestral states. (b) In the second phase, we chose ancestral states. For the root, we randomly chose a state from its possible ancestral states. For each node, we check if its parent's ancestral state is a possible ancestral state of the node. If it is, we set it as the node's ancestral state. If it is not, we set a random state from the child's possible ancestral states. In this case, a mutation occurred (+1).

**Algorithm 2** Hartigan's [3] algorithm: Build possible ancestral states

**function** PHASE1(Site $s_i$, Tree $T$)
  **traverse** $T$ **in** post-order
    **if** current node $N$ is a tip **then**
      $V(N) \leftarrow \{nucleotide(N)\}$
    **else**
      **let** $A$ and $B$ be the children of $N$.
      **if** $V(A) \bigcap V(B) \neq \emptyset$ **then**
        $V(N) \leftarrow V(A) \bigcap V(B)$
      **else**
        $V(N) \leftarrow V(A) \bigcup V(B)$
      **end if**
    **end if**
  **end traversal**
**end function**

**Algorithm 3** Hartigan's [3] algorithm: Select ancestral states

**function** PHASE2(Site $s_i$, Tree $T$)
  For the root $R$ of $T$, choose an element $S$ from $V(R)$ at random.
  **traverse** $T$ **in** pre-order ▷ Skipping the root
    **let** the current node be $A$ and its parent be $P$.
    **if** $V(P) \subseteq V(A)$ **then** ▷ We already set $V(P)$ to a single element.
      $V(A) \leftarrow V(P)$
    **else**
      $V(A) \leftarrow \{\text{RANDOMCHOICE}(V(A))\}$
    **end if**
  **end traversal**
**end function**

with always choosing the Most Significant Bit (MSB) in $\mathcal{O}(\log(16)) = \mathcal{O}(1)$ time. The overall runtime for computing the ancestral states is therefore $\mathcal{O}(n)$.

##### Encoding of the Sequences

Given a tree $T$ with $n$ sequences $S^* = \{S^1, S^2, \ldots, S^n\}$ at the tips and $n-1$ ancestral sequences $A^* = \{A^1, A^2, \ldots, A^n\}$ at the inner nodes, we can now describe the compression of a MSA (see Figure 2 and Algorithm 4). We will denote the $s$-th site of the $j$-th sequence as $S^j_s$.

To facilitate read access to random sites, we store the start of the encoding of each site in an index data structure $I$.

For each site $i$, we store the nucleotide state $s_i^{root}$ at the root sequence, followed by the changes to this site along the tree. To encode the changes, we traverse the tree in pre-order. We number the tree edges in pre-order, too. If the current node's nucleotide state for this site differs from that of its parent, we have to encode a change. We do this using the edge number leading to the current site as well as the nucleotide change mask.

**Algorithm 4** MSA compression

**function** ENCODE(Tree $T$, Sequences $S^*$) ▷ $T_{\text{EncodeTree}} + m * T_{\text{dfs}}$
  **let** $I$ be a vector mapping each site to its start location in the encoding
  ENCODETREE($T$)
  Skip space for $|S^{root}| + 1$ pointers in the output stream to later store $I$ in
  **for each** $s_i^{root} \in S^{root}$ **do**
    $I$.PUSHBACK($< i$, current position$>$)
    Write $s_i$ to output stream ▷ 4 bit
    **traverse** $T$ **in** pre-order ▷ Skipping the root
      **let** $A$ be the current node
      **let** $e_j$ be the edge from $A$'s parent to $A$; number edges by pre-order
      **if** $s_i^A \neq s_i^{parent}$ **then**
        Write <$s_i^A$, $j$>.
      **end if**
    **end traversal**
  **end for**
  $M$.PUSHBACK(`<EOF, current position + 1>`)
  Go back and write $I$
**end function**

To decode an MSA (see Figure 2 and Algorithm 5) we read the index data structure, mapping the site identifiers to the start of their encoding in the bitstream. For each site $s$ we intend to decode, we move to the specified location and start reading. The first four bits we read are the site's nucleotide state at the root sequence $s^{root}$. We then traverse the tree $T$, applying the changes along the edges. We read the changes in the same order as we wrote them, that is, in pre-order. Thus, we will never have to move backwards during the tree traversal to apply a change.

We encode sites independently of each other. We can therefore compute and write the encoding for each site sequentially on a single Processing Element (PE). Alternatively, we can distribute the sites across multiple PEs and collect the encoding afterwards. At no point in time do we have to keep all sites in memory on the same PE. We therefore do not introduce a memory bottleneck.

#### 6.2 Experimental Results

We use the algorithm described in section 6.1 to compress eleven different DNA datasets with sequences with a pairwise-identity-score ranging of 0.38 to 0.86 (see Figure 4). The compression ratio we observe ranges from

**Algorithm 5** MSA decompression

**function** DECODE(File $F$, Range of Sites $R \subseteq [1, |S^1|]$) ▷ Runtime: $|R| * T_{dfs} \in \mathcal{O}(m * n)$
  $T \leftarrow$ DECODETREE($F$)
  $I \leftarrow$ READI($F$)
  **for each** $s \in R$ **do**
    Go to start of the site's encoding in the file ▷ As indicated by $I$
    Read $S_s^{root}$ from the input stream ▷ One hot-encoded
    Read $<$ *substitutionMask*, *edgeID* $>$ from input stream
    **traverse** $T$ **in** pre-order
      Set the node's nucleotide state to its parent nucleotide state
      **if** next change is on the edge leading to the current node **then**
        Apply change-mask to the current node' state
        Read $<$ *substitutionMask*, *edgeID* $>$ from input stream
      **end if**
    **end traversal**
  **end for**
**end function**

1.9 to 21.7 with a median of 3.4 and a standard deviation of 6.5 (not normally distributed). We conclude, that the compression ratio is not sufficient to store the full MSA in memory on each node for all datasets.

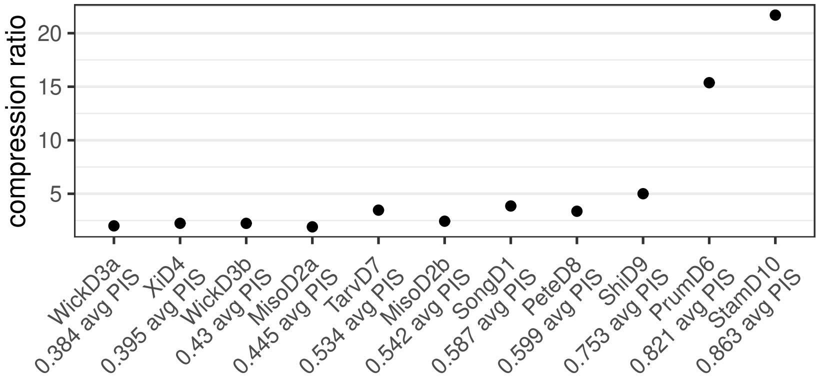


**Figure 4:** Compression ratio of different datasets using the tree-based MSA-compression. The compression ratio is defined as the size of the compressed sequences divided by the size if the uncompressed sequence. The average pairwise-sequence-identity score (avg PIS) quantifies how similar two sequences are. It is defined as the average over the number of sites each pair of sequences has in common.

[5] IUPAC. Nucleotide codes.

[6] Michael Kerrisk. Manual page of linux's `kill`. `http://man7.org/linux/man-pages/man1/kill.1.html`.

[7] Michael Kerrisk. Manual page of linux's `raise`. `http://man7.org/linux/man-pages/man3/raise.3.html`.

[8] Michael Kerrisk. Manual pages of Linux's signals. `http://man7.org/linux/man-pages/man7/signal.7.html`.

[9] Alexey M. Kozlov, Andre J. Aberer, and Alexandros Stamatakis. ExaML version 3 a tool for phylogenomic analyses on supercomputers. *Bioinformatics*, 31(15):2577–2579, March 2015.

[10] B. Misof, S. Liu, K. Meusemann, R. S. Peters, A. Donath, C. Mayer, P. B. Frandsen, J. Ware, T. Flouri, R. G. Beutel, O. Niehuis, M. Petersen, F. Izquierdo-Carrasco, T. Wappler, J. Rust, A. J. Aberer, U. Aspock, H. Aspock, D. Bartel, A. Blanke, S. Berger, A. Bohm, T. R. Buckley, B. Calcott, J. Chen, F. Friedrich, M. Fukui, M. Fujita, C. Greve, P. Grobe, S. Gu, Y. Huang, L. S. Jermiin, A. Y. Kawahara, L. Krogmann, M. Kubiak, R. Lanfear, H. Letsch, Y. Li, Z. Li, J. Li, H. Lu, R. Machida, Y. Mashimo, P. Kapli, D. D. McKenna, G. Meng, Y. Nakagaki, J. L. Navarrete-Heredia, M. Ott, Y. Ou, G. Pass, L. Podsiadlowski, H. Pohl, B. M. von Reumont, K. Schutte, K. Sekiya, S. Shimizu, A. Slipinski, A. Stamatakis, W. Song, X. Su, N. U. Szucsich, M. Tan, X. Tan, M. Tang, J. Tang, G. Timelthaler, S. Tomizuka, M. Trautwein, X. Tong, T. Uchifune, M. G. Walzl, B. M. Wiegmann, J. Wilbrandt, B. Wipfler, T. K. F. Wong, Q. Wu, G. Wu, Y. Xie, S. Yang, Q. Yang, D. K. Yeates, K. Yoshizawa, Q. Zhang, R. Zhang, W. Zhang, Y. Zhang, J. Zhao, C. Zhou, L. Zhou, T. Ziesmann, S. Zou, Y. Li, X. Xu, Y. Zhang, H. Yang, J. Wang, J. Wang, K. M. Kjer, and X. Zhou. Phylogenomics resolves the timing and pattern of insect evolution. *Science*, 346(6210):763–767, November 2014.

[16] Alexandros Stamatakis, Markus Göker, and Guido W. Grimm. Maximum likelihood analyses of 3,490 rbcl sequences: Scalability of comprehensive inference versus group-specific taxon sampling. *Evolutionary Bioinformatics*, 6:EBO.S4528, 2010. PMID: 20535232.

[17] Steinbruch Center for Computing (SCC). ForHLR - hardware and architecture. `https://wiki.scc.kit.edu/hpc/index.php/ForHLR_-_Hardware_and_Architecture`, 2020.

[18] Steinbruch Center for Computing (SCC). Konfiguration des ForHLR II. `https://www.scc.kit.edu/dienste/forhlr2.php`, 2020.

[19] James E. Tarver, Mario dos Reis, Siavash Mirarab, Raymond J. Moran, Sean Parker, Joseph E. O'Reilly, Benjamin L. King, Mary J. O'Connell, Robert J. Asher, Tandy Warnow, Kevin J. Peterson, Philip C. J. Donoghue, and Davide Pisani. The interrelationships of placental mammals and the limits of phylogenetic inference. *Genome Biology and Evolution*, 8(2):330–344, January 2016.

[20] Norman J. Wickett, Siavash Mirarab, Nam Nguyen, Tandy Warnow, Eric Carpenter, Naim Matasci, Saravanaraj Ayyampalayam, Michael S. Barker, J. Gordon Burleigh, Matthew A. Gitzendanner, Brad R. Ruhfel, Eric Wafula, Joshua P. Der, Sean W. Graham, Sarah Mathews, Michael Melkonian, Douglas E. Soltis, Pamela S. Soltis, Nicholas W. Miles, Carl J. Rothfels, Lisa Pokorny, A. Jonathan Shaw, Lisa De Gironimo, Dennis W. Stevenson, Barbara Surek, Juan Carlos Villarreal, Béatrice Roure, Hervé Philippe, Claude W. De Pamphilis, Tao Chen, Michael K. Deyholos, Regina S. Baucom, Toni M. Kutchan, Megan M. Augustin, Jun Wang, Yong Zhang, Zhijian Tian, Zhixiang Yan, Xiaolei Wu, Xiao Sun, Gane Ka Shu Wong, and James Leebens-Mack. Phylotranscriptomic analysis of the origin and early diversification of land plants. *Proceedings of the National Academy of Sciences of the United States of America*, 111(45):E4859–E4868, November 2014.

[21] Zhenxiang Xi, Liang Liu, Joshua S. Rest, and Charles C. Davis. Coalescent versus concatenation methods and the placement of amborella as sister to water lilies. *Systematic Biology*, 63(6):919–932, July 2014.

[22] Ya Yang, Michael J. Moore, Samuel F. Brockington, Douglas E. Soltis, Gane Ka-Shu Wong, Eric J. Carpenter, Yong Zhang, Li Chen, Zhixiang Yan, Yinlong Xie, Rowan F. Sage, Sarah Covshoff, Julian M. Hibberd, Matthew N. Nelson, and Stephen A. Smith. Dissecting molecular evolution in the highly diverse plant clade caryophyllales using transcriptome sequencing. *Molecular Biology and Evolution*, 32(8):2001–2014, April 2015.
